# Simultaneous Identification of Brain Cell Type and Lineage via Single Cell RNA Sequencing

**DOI:** 10.1101/2020.12.31.425016

**Authors:** Donovan J. Anderson, Florian M. Pauler, Aaron McKenna, Jay Shendure, Simon Hippenmeyer, Marshall S. Horwitz

## Abstract

Acquired mutations are sufficiently frequent such that the genome of a single cell offers a record of its history of cell divisions. Among more common somatic genomic alterations are loss of heterozygosity (LOH). Large LOH events are potentially detectable in single cell RNA sequencing (scRNA-seq) datasets as tracts of monoallelic expression for constitutionally heterozygous single nucleotide variants (SNVs) located among contiguous genes. We identified runs of monoallelic expression, consistent with LOH, uniquely distributed throughout the genome in single cell brain cortex transcriptomes of F1 hybrids involving different inbred mouse strains. We then phylogenetically reconstructed single cell lineages and simultaneously identified cell types by corresponding gene expression patterns. Our results are consistent with progenitor cells giving rise to multiple cortical cell types through stereotyped expansion and distinct waves of neurogenesis. Compared to engineered recording systems, LOH events accumulate throughout the genome and across the lifetime of an organism, affording tremendous capacity for encoding lineage information and increasing resolution for later cell divisions. This approach can conceivably be computationally incorporated into scRNA-seq analysis and may be useful for organisms where genetic engineering is prohibitive, such as humans.

## INTRODUCTION

A fundamental question of developmental biology addresses how the fertilized egg, through a repertoire limited to cell division, migration, differentiation, and death, matures into a multicellular organism. Sulston and colleagues employed a microscope, pen, and paper to painstakingly record the history of each of the 959 cells of the transparent nematode *C. elegans* during its 14-hour development (Sulston et al., 1983). However, cell fate determination in vertebrates remains daunting because there are vastly more cells, more types of cells, more time required for development, opaque tissues, and, unlike *C. elegans*, lineage is not invariant between individuals.

Other experimental approaches have utilized embryonic chimerism and cell marking with dyes (Salipante and Horwitz, 2007), as well as transgenic markers and barcodes to track clonal histories (Weissman and Pan, 2015). However, the relationship between the number of clonally related cells and permissible lineages grows explosively. For example, for a 32-cell embryo, more than 10^42^ distinct lineage histories are possible (Salipante and Horwitz, 2007). A massive capacity for uniquely labeling cells is therefore required to unambiguously resolve lineage.

In recent years, single cell genomic and transcriptomic analyses have accelerated capabilities for decoding cell lineage and identity (Wagner and Klein, 2020).

One approach for deciphering cell lineage is to retrospectively infer the sequence of mutations acquired in single cells (Carlson et al., 2011; Frumkin et al., 2008; Ju et al., 2017; Lee-Six et al., 2018; Lodato et al., 2015; Ludwig et al., 2019; Salipante and Horwitz, 2006). Mutations are sufficiently frequent such that a large proportion of cells acquire a genome unique unto themselves. In fact, brute force sequencing of single cell genomes provides sufficient information to unambiguously infer the lineage of mouse cells, tracing back to the zygote, albeit at low resolution due to the infrequency of spontaneous mutations (Behjati et al., 2014).

More recently, engineered recorder systems, such as GESTALT (McKenna et al., 2016) and MEMOIR (Frieda et al., 2017), which induce mutations with CRISPR or other means to generate cellular barcodes, have enabled high throughput lineage reconstruction from zebrafish and other model organisms (Wagner and Klein, 2020).

At the same time, breakthroughs in single cell RNA sequencing (scRNA-seq), in combination with algorithmic advances, have allowed for elucidation of cell state trajectories (Packer and Trapnell, 2018; Trapnell et al., 2014). However, the relationship between a cell’s developmental trajectory, as reflected in its transcriptome, and its lineage, as defined by its history of cell divisions, remains an important and challenging question (Wagner and Klein, 2020). Engineered recorder systems can extract both cell state and lineage at single cell resolution by incorporating barcodes into a transcript that can be read out through scRNA-seq (Raj et al., 2018; Wagner and Klein, 2020). Of course, genetic engineering cannot be used to study development in humans or other organisms not suitable for genetic engineering. Previous approaches in humans for correlating cell state with lineage based on somatic mutations have required parallel RNA and DNA sequencing, respectively (Huang et al., 2020).

In principle, somatic mutations occurring within genes should also be detectable in the transcriptome. Mutational analysis of transcripts solves a principal technical challenge for detecting mutations in single cells. In order to interrogate multiple markers or perform sequencing on a single genome, whole genome amplification (Frumkin et al., 2008), which is error-prone (Sabina and Leamon, 2015), or *in vitro* culture of clonally expanded cells (Behjati et al., 2014; Salipante et al., 2008), which is difficult to scale, is required. In contrast, thousands of copies of a given RNA molecule may be present in a single cell. However, the transcriptome comprises only a fraction of the genome, and mutations concentrate in hotspots (Alexandrov et al., 2013), typically non-transcribed repetitive sequences, which are difficult to sequence (Shendure and Ji, 2008), even if transcribed. Accuracy limitations impose additional challenges for detecting mutations through RNA sequencing: RNA polymerase has an error rate of ~10^−5^, which is about 5,000-fold higher than point mutation frequency (Gout et al., 2017), apparent mutations often reflect RNA edits (Ding et al., 2019), reverse transcriptase used to generate cDNA sequencing libraries has a fidelity of ~10^−4^ (Ji and Loeb, 1992), and next-generation sequencing misreads ~10^−3^ bases (Minoche et al., 2011).

Detection of loss of heterozygosity (LOH) offers a potential workaround. While somatic point mutations are not uncommon, LOH, which can arise from somatic recombination or other mechanisms, occurs frequently. The phenomenon was first described as ‘twin spotting’ in *Drosophila*, more than 80 years ago (Stern, 1936). LOH can result in surprisingly frequent reversion of germline disease mutations (Revy et al., 2019). For example, in ichthyosis with “confetti,” innumerable revertant clones densely speckle skin (Choate et al., 2010). Measurements at marker genes (LaFave and Sekelsky, 2009; Larson et al., 2006; Moynahan and Jasin, 2010) indicate that LOH occurs with a frequency of ~10^−4^−10^−5^/locus/cell division. Multiple regions of LOH of variable length are distributed throughout the genome, thereby potentially uniquely marking each cell.

Compared with the identification of somatic point mutations, detection of LOH offers advantages: LOH occurs frequently and affords voluminous informational coding capacity. LOH represents a homogenization of tolerated genetic variants, less likely to skew cell growth patterns. LOH can be assayed at sites known to be germline heterozygous and are therefore predictable and dramatically more economical to survey than mutations, which can arise anywhere throughout the genome. The signal of LOH will be reinforced by multiple adjacent informative loci spanning large chromosomal segments. LOH corresponds to heterozygous base positions converting to homozygosity, whereas most sequencing errors involve false calls of heterozygosity (Shendure and Ji, 2008).

LOH distributed throughout the genome may be identifiable in the transcriptome as tracts of apparent monoallelic expression in contiguous genes. However, not every gene is expressed in all tissues nor at the same time or levels, scRNA-seq sampling can be sparse, and the phenomenon would go undetected in inbred genetic backgrounds, since there is no heterozygosity to be lost.

Here we investigate a computational strategy for overcoming these challenges and employ scRNA-seq to identify cell state, based on gene expression, while simultaneously extracting cell lineage, based on detection of LOH manifesting as tracts of monoallelic expression.

In an initial application, we have studied major cellular classes during development of mouse cerebral cortex at single cell resolution.

## RESULTS

### Cortical Cell Identification

We analyzed single cells from the cerebral cortex of eight mice (**Fig. 1A**). In order to distinguish parental alleles, we studied F1 offspring of a cross between two different inbred strains, C57Bl/6J (B6) and CAST/EiJ (CA), whose genomes have previously been sequenced. To control for parental sex, four mice were products of matings of female B6 and male CA parents, and four mice were born to parents in which strain sexes were reversed. From each group of four, we analyzed a female and male at two developmental time points, postnatal days 0 (P0) and 42 (P42). We focused our analysis on *Emx1*^*+*^ cortical projection neuron and glia lineage. To this end, we utilized B6 mice transgenic for *Emx1*-Cre;*Z/EG*, such that enhanced green fluorescent protein was expressed in cells that, at least at some point during their development, had expressed the neuronal transcription factor EMX1 (Gorski et al., 2002). *Emx1*-marked cells were isolated by flow cytometry and underwent scRNA-seq to a high depth of coverage using Smart-seq2 (Picelli et al., 2013). Approximately 50 cells from each mouse cortex (404, in total) passed quality control filtering (median of 1,735,775 unique reads per cell).

**Figure 1.**
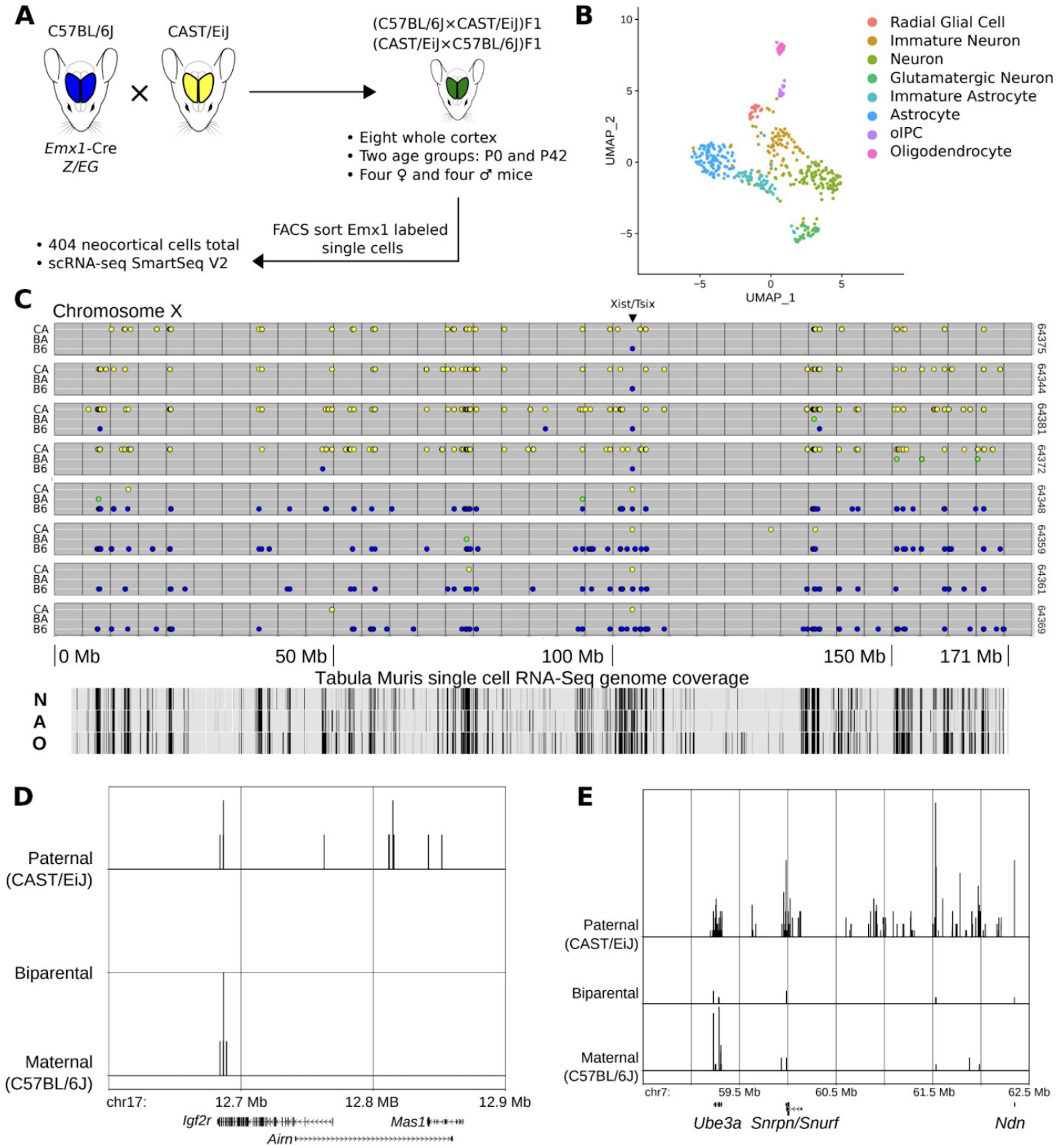
Isolation and whole transcriptome sequencing of mouse neocortical cells. (A) B6 (blue) and CA (yellow) mice were crossed in both directions to create heterozygous F1 offspring (green). Single *Emx1* expression-marked neocortical cells were isolated from two different ages and their transcriptomes sequenced. (B) 404 cells were clustered in dimensionally reduced (UMAP) space based on gene expression, and eight cell types were identified. oIPC = oligodendrocytic intermediate progenitor cell. (C) Representative allele plots of heterozygous SNV loci detected along the X chromosome in 8 cells (rows) from a female P0 mouse showing X-inactivation. Note the reciprocal allele state detected at the *Xist/Tsix* locus. Allele state: blue (B6) = B6, green (BA) = B6:CA, yellow (CA) = CA. N = neuron, A = astrocyte, O = oligodendrocyte. (D & E) Relative density histogram ridgeline plots (1 bp bins) of cells expressing maternal, paternal, or biparental variants at particular SNV locations from one mouse (P0-2, 64 cells). Chromosome coordinates and relevant genes are indicated on the x-axis. Density for each allele category is indicated on the y-axis. (D) *Igf2r* locus. (E) Prader-Willi/Angelman syndrome locus.

Using known markers of cell type (**Supplemental Fig. 1** and **Supplemental Table 1**), scRNA-seq analysis of cells from all mice combined led to identification of eight cell types (**Fig 1B**).

Radial glial cells (RGC) are neural progenitors defined by expression of several genes marking transition from neuroepithelial to mesenchymal states. We used expression of the genes encoding intermediate filaments Vimentin (*Vim*) and Nestin (*Nes*), the extracellular matrix component Tenascin C (*Tnc*), neurogenic transcription factors *Pax6 and Sox2*, as well as glial markers GLAST (*Slc1a3*) and Blbp (*Fabp7*) to identify these cells.

We defined three classes of neurons using a variety of markers. Doublecortin (*Dcx*) and *Neurod1* expression arises in the immature neural cluster and decreases in the more mature neuron and glutamatergic neuron clusters. Beta III Tubulin (*Tubb3*) transcription increases in the immature cluster and peaks in the neuron cluster. The immature neuron cluster also shows marked increase in *Sema3c* and *Sox11* transcription, suggesting that these cells are radially migrating (Hoshiba et al., 2016; Wiegreffe et al., 2015), while the neuronal cluster is post-migratory. The post-mitotic neuronal marker NeuN (*Rbfox3*) shows low expression in immature neuron and neuron clusters, with a marked increase in the glutamatergic neuron cluster. Neuronal cytoskeleton components *Nefm* and *Nefh* also demonstrate increased transcription in glutamatergic neurons. The glutamatergic cluster is marked by the vesicular glutamate transporter vGluT1 (*Slc17a7*), the NMDA receptor subunit of the glutamate receptor channel GluN1 (*Grin1*), and glutaminase (*Gls*). Together these results describe the general maturation of a neuron to a functional state and are expected, as *Emx1* expression marks excitatory pyramidal projection neurons in the mouse cortex (Gorski et al., 2002).

Astrocytes, in general, are distinguished by the presence of GLAST, the glutamate/aspartate transporter Glt-1 (*Slc1a2*), and glutamine synthetase (*Glul*). We also identified a separate cluster of astrocytes that are marked by increased expression of the calcium binding component *S100b* and the brain-specific aldolase C (*Aldoc*). *S100b* expression marks commencement of astrocyte terminal differentiation (Raponi et al., 2007).

Two oligodendrocyte populations were identified. The first, corresponding to an oligodendrocytic intermediate progenitor population (oIPC), is uniquely marked by platelet-derived growth factor receptor *Pdgfra* and proteoglycan Ng2 (*Cspg4*). Expression of the transcription factor *Sox10* increases in these cells while continuing at a lower level in oligodendrocytes. The second mature oligodendrocyte population is identified by increased expression of the tight junction component Claudin 11 (*Cldn11*) and Myelin Oligodendrocyte Glycoprotein (*Mog*).

In general, the identified cell types were consistent with the age of sampled cortex. Analysis of cell cycle-related genes (**Supplemental Table 2**) revealed that clusters identified as mature and predicted to be post-mitotic are predominantly classified as G1 phase cells (**Supplemental Fig. 2**). Most P42 cells are classified as G1, while the oIPC of those mice exhibit populations in both S and G2/M phases. RGC, immature neurons, and neurons display more heterogeneous distributions within the cell cycle This is expected for RGC, but not for neurons, which are post-mitotic. However, our findings are consistent with previous reports suggesting that, on a transcriptional level, the progression to a post-mitotic state in neurons is gradual and that cell cycle genes are expressed but do not lead to productive cell division (Anda et al., 2016).

### Detection of Allele-Specific Expression

A cross between different inbred mouse strains, each homozygous throughout their diploid genome, should predictably generate heterozygosity in F1 progeny wherever parental strains differ in DNA sequence, consisting of indels, microsatellite polymorphisms, and single nucleotide variants (SNVs). Of these, SNVs are most readily interrogated by scRNA-seq. Using the published genomes of the B6 and CA parental strains, we constructed a list of 20,667,142 SNVs across all autosomes and the X chromosome, which serves as a guide for determining expressed allele status in each cell.

For further studies, we focused on the four mice (one of each sex, for P0 and P42 stages) derived from a female B6 crossed to a male CA. For each mouse, we interrogated a mean of 200,540 (standard deviation 67,959) informative heterozygous SNVs distributed across mapped scRNA-seq reads in our dataset, corresponding to an average density of 81 loci per megabase (Mb). SNV positions that were homozygous for either parental variant suggest monoallelic expression, potentially consistent with LOH, while heterozygosity would indicate that both parental alleles are present, excluding the possibility of LOH. When using scRNA-seq information as a starting point, the number of loci that yield allele information is dependent on cell type and its corresponding transcriptome. Recognizing that not all cells express the same transcript, as well as incomplete capture of all transcripts, the median number of transcribed SNV coordinates that passed quality filters for any given cell for each mouse was 10,805 (LQ 7,232; UQ 14,160), yielding an average median autosomal coverage density of 4 SNVs/Mb (LQ 3; UQ 6). These results show that scRNA-seq generated transcriptomes have the potential to provide allele state information at a density comparable to commercial DNA-based genomic SNV microarrays clinically employed for detection of LOH and copy number variants in the ~20% larger human genome (e.g., Affymetrix High Density, 750K loci; Agilent Medium Density, 30K loci; Oxford Gene Low Density, 6,186 loci), although we do not explore here whether scRNA-seq data are useful for detecting copy number changes.

### Detection of X-Inactivation and Imprinting with scRNA-seq

A measure of our variant calls and filtering methods can be provided by examining regions with predictable allele-specific expression patterns. Most obvious is the X-chromosome in female mice. One of the two X-chromosomes is randomly inactivated during embryogenesis in female mice, leading to expression from predominantly one chromosome or the other (Galupa and Heard, 2018). Certain regions on the inactive chromosome, however, escape inactivation, and, in fact, some are responsible for maintaining the inactive state. For the purpose of our studies, these exceptions offer an important test of whether biallelic expression can be identified at loci that might otherwise be interpreted as monoallelic.

Plots of the X-chromosome from eight representative cells of a B6 (female) × CA (male) F1 P0 female hybrid mouse are shown in **Fig. 1C**. For each cell, allele frequencies of informative SNVs are plotted along the length of the chromosome. Their position on the vertical axis, along with color, indicates which parental allele(s) are detected, with monoallelic CA (yellow) shown on top, monoallelic B6 (blue) on the bottom, and biparental expression (green) in-between. Four cells exhibit inactivation of the B6 X-chromosome, while the other four cells reveal inactivation of the CA X-chromosome, as indicated by the predominant expression of the opposite parental allele. A reversal of this pattern occurs predictably at the *Xist/Tsix* locus. *Xist* and *Tsix* are exclusively expressed, respectively, from the inactive and active X-chromosomes and contribute to X-inactivation (Galupa and Heard, 2018). Some cells show regions with inconsistent escape from X-inactivation, including genes previously observed to exhibit leaky expression (**Supplemental Table 3**). Complicating matters is that detection of escape from X-inactivation in hybrid mice appears dependent on parental strains and can be tissue specific (Andergassen et al., 2017; Berletch et al., 2015). Nevertheless, for the X-chromosome, our data capture the expected variation in allele-specific transcription, with each cell displaying a dominant parental variant in female mice.

A few autosomal segments exhibit imprinting, containing genes in which only one or the other parental allele is expressed, thereby offering another opportunity to assess accuracy for identifying loci with biallelic expression amidst regions of allele-specific expression. A cluster of imprinted genes on mouse chromosome 17 includes insulin-like growth factor II receptor (*Igf2r*) and lncRNA *Airn* (Latos et al., 2012). Unlike humans, mouse *Igf2r* is exclusively maternally expressed in most peripheral tissues. However, *Igf2r* imprinting is relaxed in mouse astrocytes and oligodendrocytes, giving a more human-like biparental expression pattern (Hu et al., 1998). We detect cells expressing either maternal or paternal *Igf2r* alleles, along with exclusively paternal expression of *Airn* (**Fig. 1D**).

Another imprinted locus is the Prader-Willi/Angelman syndrome (PWS/AS) region of mouse chromosome 7, which consists of several imprinted genes, expressing only paternal copies of *Snrpn*, *Snurf*, *Ipw*, and *Npn*, along with exclusively maternal expression of *Ube3a* (Bervini and Herzog, 2013). Developing neurons and glial cells, however, biallelically express *Ube3a (Judson et al., 2014)*. We again capture this relaxation of imprinting (**Fig. 1E**). As expected, expression at other loci (*Snrpn*, *Snurf*, and *Npn*) is predominantly paternal, though we also observe low levels of biallelic or maternal expression.

The examples above demonstrate that our variant calling pipeline and quality filters are sufficient to produce positive and specific biallelic variant calls in well-studied regions of the genome where monoallelic expression is known to occur, thus mitigating false inference of LOH due to epigenetic factors. In addition, we can capture predicted tissue dependent allele expression patterns. These results suggest that these data are of sufficient quality to detect regions of LOH with specificity in single cells.

### Inferring Loss of Heterozygosity in Single Cells

Compared to bulk RNA-seq, scRNA-seq may artifactually indicate monoallelic expression due to stochastic sampling biases, especially for genes with modest levels of expression, and transcriptional “bursting,” in which, at the time sequencing is performed, only one active allele is captured (Borel et al., 2015; Deng et al., 2014; Finn and Misteli, 2019; Reinius and Sandberg, 2015; Reinius et al., 2016). The latter concern is at least partly mitigated by the fact that transcriptional bursts are typically shorter than the half-lives of RNA (Finn and Misteli, 2019). Nevertheless, these phenomena complicate interpretation of allele states. The chance that an apparently monoallelic SNV reflects sampling noise due to the nature of the scRNA-seq protocol, rather than an underlying LOH event, must therefore be considered. To take advantage of the fact that contiguous tracts of monoallelic SNVs help validate interpretation of reads at adjacent positions, we employed a hidden Markov model (HMM) to infer the most likely genotype corresponding to observed patterns of allele specific expression.

We posit three hidden states relating to a cell’s genotype: heterozygous, homozygous B6, or homozygous CA. A transition from the heterozygous state to either homozygous state is 10,000-fold less likely than remaining in the heterozygous state, based on observed frequencies of interhomolog chromosomal exchanges (Larson et al., 2006). Within a stretch of homozygosity created by recombination, transitioning back to the heterozygous state would require a second recombination event, which observations suggest is 10-fold less likely than a single event (i.e., overall 100,000-fold less likely than remaining in the particular state) (Larson et al., 2006). Transitions from one homozygous genotype to the other cannot be generated through recombination. The ultimate effect of these transition rates is that, given an observed homozygous run of either allele, there is a small bias for continuation of calling a homozygous state, instead of transitioning back and forth between heterozygous and homozygous states.

The probability of observing a particular SNV in the heterozygous state or either homozygous state is described by the emission state matrix. Due to both bursting and incomplete sampling, we estimated that biallelic SNVs will be observed correctly as heterozygous in the scRNA-seq data with a probability of 0.10 but will be more frequently incorrectly observed as monoallelic for the maternal allele with probability 0.45 or the paternal allele also with a probability of 0.45, as was demonstrated by comparing bulk to single cell sequencing on the same sample (Borel et al., 2015). A truly homozygous locus should only be observed as monoallelic. An observation of either heterozygous SNVs or homozygosity for a SNV from the opposite allele would switch the hidden state back to heterozygosity.

In our model, the heterozygous to homozygous transition probability only modestly influences determination of tracts of LOH. A ten-fold decrease in probability (i.e., from 10^−4^ to 10^−5^) results in a requirement for just three more continuously observed monoallelic SNVs necessary to assign a region of LOH (**Supplemental Fig. 3**). We therefore selected an intermediate transition probability to account for genome wide variance in observed frequencies of LOH (Larson et al. 2006). In fact, most LOH events would still be called using this model if the transition probability were set to 10^−8^.

We used a Viterbi path, which describes the most likely set of hidden states and transitions given an observed dataset, to infer underlying genotypes for the observed readout across autosomal SNVs from scRNA-seq data. Examples of chromosome 19 from two cells, one with evidence of LOH and one without, are shown in **Fig. 2A**. For each cell, two plots are shown. The upper plot illustrates the observed allele state for any SNV based on scRNA-seq data, while the bottom plot shows the most likely HMM-inferred genotype. Note, as well, that the density of SNV coverage corresponds to which genes are expressed in each cell type, as shown by transcripts mapped for relevant cell types in the Tabula Muris project (Tabula Muris Consortium et al., 2018). Cell 64461 demonstrates the tendency for scRNA-seq to capture SNVs within transcripts corresponding to only one allele. Most SNVs are observed as homozygous for either parental allele (colored yellow or blue by parent of origin). Nevertheless, biallelically expressed SNVs (green) and monoallelically expressed SNVs from either parent are interspersed throughout. Consequently, the inferred genotype for this chromosome is heterozygous, with no LOH events, throughout its length. This interpretation is consistent with previous studies showing that *in silico* aggregation of what may appear to be predominately monoallelic scRNA-seq data becomes biallelic, matching bulk RNA-seq data from another sample of the same cell line (Borel et al., 2015). The second cell (64474) shows a similar stochastic monoallelic expression pattern, but two LOH events are inferred. The first is an interstitial event proximal to the centromere while the second is a more telomeric interstitial event.

**Figure 2.**
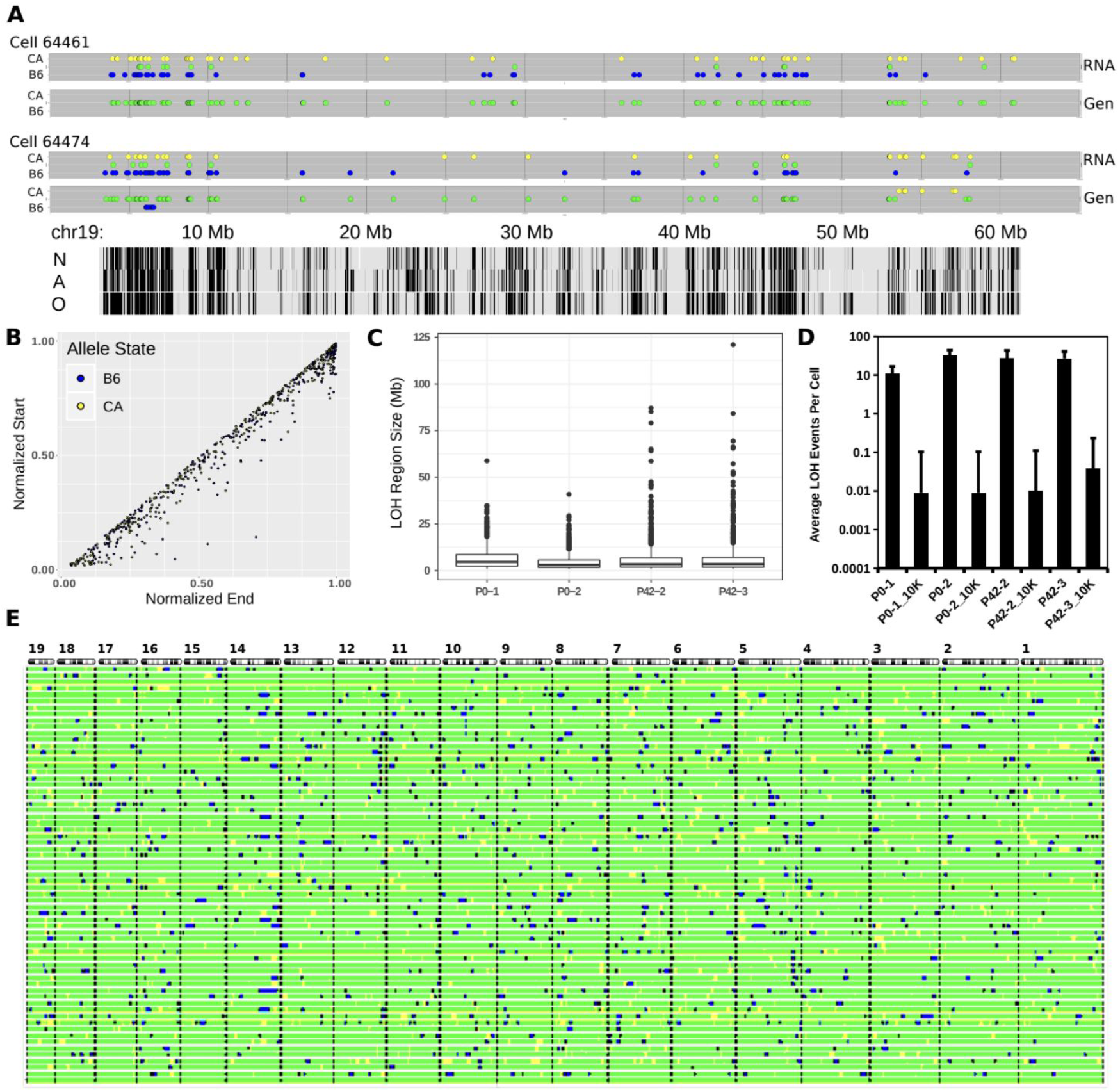
Inferring loss of heterozygosity regions from scRNA-seq. (A) Hidden Markov Model (HMM) results showing two cells with either no LOH event (cell 64461) or two events (cell 64474). Detected loci are depicted as a plot for chromosome 19 with the horizontal axis indicating chromosome position and the vertical axis indicating expressed allele state (RNA) or HMM inferred genotype (Gen). For each cell, the scRNA-seq data is shown on top with the most likely HMM genotype shown below. Tabula Muris cell-type specific transcription tracks are shown at the bottom. N = neuron, A = astrocyte, O = oligodendrocyte. (B) Normalized chromosome positions of all LOH events from all autosomes of one mouse (P0-1). (C) Distribution of LOH lengths for all four mice showing median (bar) and IQR (box). (D) Average LOH events per cell for each mouse and the average LOH events of 10,000 randomly sampled *in silico* cells derived from the respective mouse. (E) Autosomal barcode of 56 cells plus a virtual zygote (bottom bar) derived from one mouse (P0-2). Autosomes are shown at the top with the centromeres on the left. Black bars indicate autosomal boundaries. For all panels: blue = B6, green = B6:CA, yellow = CA.

Applying this approach to all cells across all mice, we observe predominantly interstitial LOH events (**Fig. 2B**). For all called LOH regions across all autosomes and cells, mean, standard deviation, and number of events are as follows: P0-1 - 5.30, 6.14, 795; P0-2 - 2.80, 3.65, 3447; P42-2 - 3.69, 6.44, 2469; P42-3 - 3.44, 6.93, 2381.

Consistent with this, data from high density SNP microarrays show that the predominant LOH event in healthy tissue is interstitial (Melcher et al., 2011; Mohamedali et al., 2007; O’Keefe et al., 2010). The size distribution of LOH events we detect (**Fig. 2C**) is also similar to reported values. For example, for the P0-1 mouse, we find a median size of 4.6 Mb, (range 1.0 - 29 Mb), compared to median estimates (in humans) of 1.2 Mb (range 0.3 - 6.7 Mb) (Mohamedali et al., 2007) and 8.7 Mb (range 0.3 - 65 Mb) (O’Keefe et al., 2010). In sum, our method of detecting LOH from scRNA-seq information appears to capture LOH events similar in size and chromosomal position to those observed using high density SNP microarrays based on genomic DNA.

As noted, there is a tendency for scRNA-seq data, and in particular those that don’t utilize unique molecular identifiers (UMIs), to incorrectly imply monoallelic expression at any particular locus. To determine whether or not LOH events detected by our algorithm reflect noise or other sampling issues, we created 10,000 *in silico* “cells” for each mouse by scrambling the collected scRNA-seq data. Briefly, for each SNV in a single cell in any one mouse, we sampled, with replacement, from the entire mouse cell set whether or not that locus was expressed and, if so, its scRNA-seq state (B6, B6:CA, or CA). This was performed for all detected autosomal loci, and the results were stored for that particular cell. This process was repeated 10,000 times, and each cell was then processed in the same manner as our original single cells. The results are shown in **Fig. 2D**. For all four mice the average number of LOH events per cell determined from the actual data is ~15, while the average number of LOH events per cell for randomized data is ~0.02. An approximately 1,000-fold reduction in LOH detection in randomized data indicates that our detected events are unlikely to represent an scRNA-seq sampling artifact.

The allelic ratio for SNVs distinguishing parental origin of expressed genes has been shown to correlate in scRNA-seq datasets for distances of up to about 500 kb (Borel et al., 2015). This correlation is due in most part to SNVs locating within the same gene, and hence being contained within the same sequenced RNA molecule. Correlation between SNVs residing in two separate yet coordinately transcribed genes could conceivably also be observed when the genes reside within the same topologically associating domain, which are similarly between about 200 kb and 1 Mb in length (Finn and Misteli, 2019). To control for such phenomena, we excluded regions of monoallelic expression shorter than 1 Mb from our analysis. A filter of this size also minimizes inclusion of loci that may be physiologically imprinted, as noted above, or otherwise monoallelically expressed, such as olfactory receptors or immunoglobulins (Khamlichi and Feil, 2018).

SNVs are neither uniformly distributed across a chromosome nor are the transcripts in which they are located expressed in all cells. This creates ambiguity in precisely defining the beginning and ending of LOH regions. We consequently assigned beginning and ending points for each region to 2 Mb bins tiled across the length of each chromosome. Using these conditions, we created an autosomal map of LOH events for each cell. This map, shown in **Fig. 2E**, represents a cellular barcode uniquely identifying each cell. The collection of LOH events represent heritable somatic mutations that, just like single nucleotide, microsatellite, or CRISPR-Cas9 induced variants, can be used to infer cell lineage.

The distribution of LOH events shared between cells within each mouse is shown in **Supplemental Fig. 4A**. Most LOH events are unique to a single cell, but as many as nine are shared across cells within a particular mouse. Boundaries for LOH events are only approximated as a result of the fact that not all transcripts are expressed in all cells. Nevertheless, most LOH events appear unique to each mouse (**Supplemental Fig. 4B**).

To further address whether an LOH event is more likely to occur in cells from the same mouse compared to a different mouse, we devised an enrichment quotient. For each LOH event, the numerator represents the number of cells containing that allele for one of the four mice, and the denominator is the sum of all occurrences of that allele across all cells from that mouse and one other mouse. A value of one would indicate that the allele is unique with respect to the particular “target” mouse, whereas a value of ~0.5 would indicate that the allele is just as likely to be found in cells from the other mouse it is compared to (approximate because of differences in the number of LOH events for each mouse). We report the mean quotient for all alleles for a particular mouse (**Supplemental Table 4**). For each unidirectional pairwise comparison, the enrichment quotient ranges from 0.82-0.96, meaning that, on average, inferred LOH events tend to recur within cells from the same mouse.

### Loss of Heterozygosity as a Marker of Lineage

To deduce cellular lineage, we employed Camin-Sokal parsimony, in which the most likely phylogeny minimizes the number of ordered character state changes from the base of the dendrogram to its tips (Camin and Sokal, 1965). LOH events for any one chromosome can be described as a discrete and irreversible event. We coded autosomal LOH events occurring on the same chromosome as a series of two-state characters, with heterozygosity being the ancestral character. In F1 mice, LOH events should not exist in the zygote. We implemented this model in a Bayesian phylogenetic analysis using the MrBayes algorithm (Ronquist et al., 2012). For each mouse we added a cell representing the zygote, with no LOH events assigned. The zygote branch length is expected to be close to zero and placed as the outgroup in the resultant dendrogram. We excluded CNVs on the sex chromosomes.

The consensus phylogram for a P0 (P0-2) mouse is shown in **Fig. 3A**. The zygote occurs as an outgroup when compared to the rest of the cells (with an exception for just one other cell) and shows the least amount of change (less than 1 LOH event), as expected. Most changes occur towards the tips of the tree, and intranodal distances are short. Pairwise comparison of differences between cells exhibits a unimodal distribution (**Supplemental Fig. 5**). Such a pattern, along with short intranodal distances, is consistent with expanding populations, as expected during embryogenesis; in contrast, populations of constant size exhibit more evenly spaced intranodal distances and monotonically decreasing pairwise-distance distributions (Rogers and Harpending, 1992). Given these results, we believe that our evolutionary model reflects the sequence of LOH events acquired during development.

**Figure 3.**
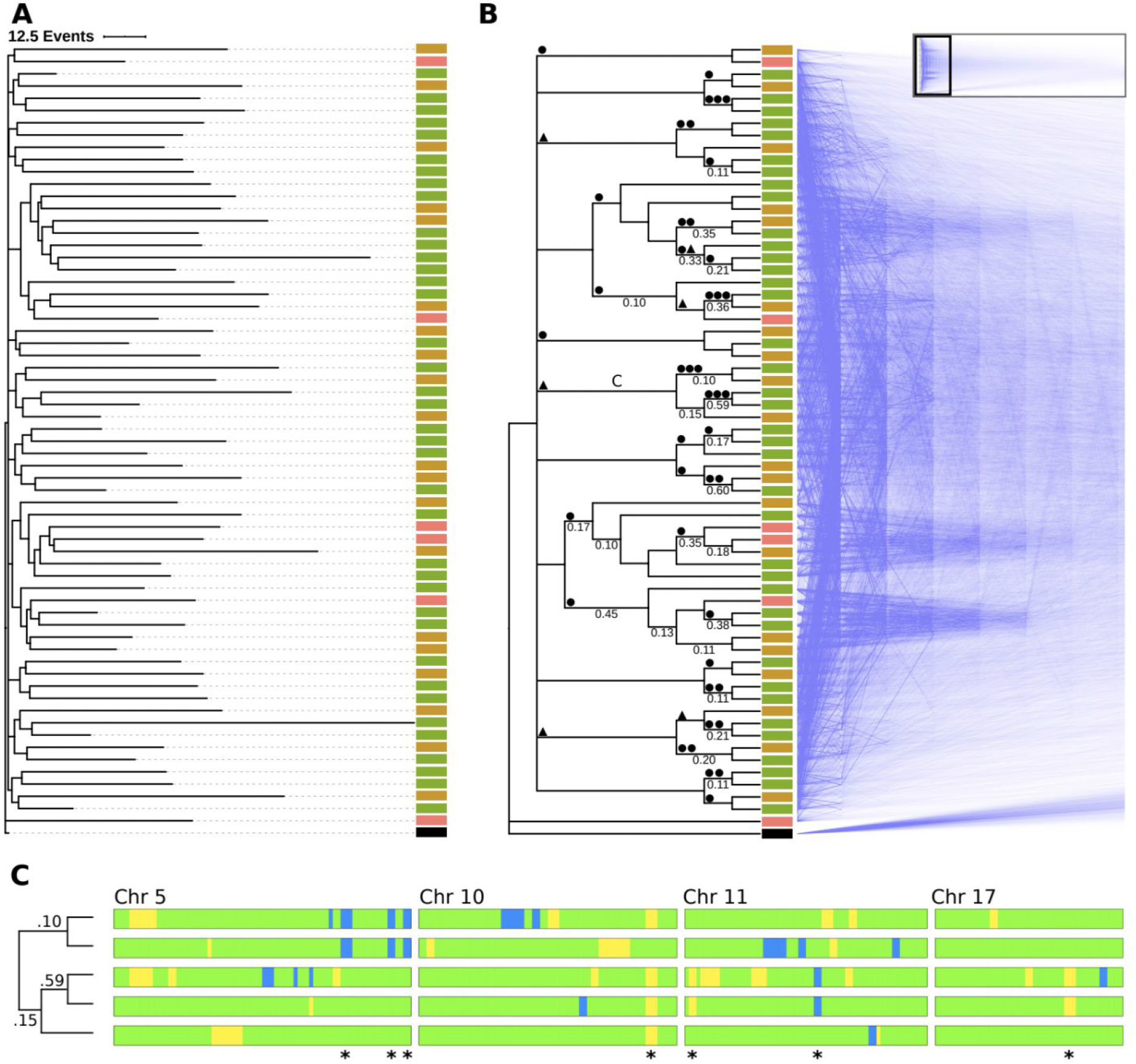
Phylogenetic analysis using LOH events. The lineage of 56 cortical cells plus a virtual zygote from a P0 mouse was inferred using a Camin-Sokal parsimony-inspired Bayes model. (A) A consensus phylogram showing the relatedness and number of LOH events for each cell. Scale bar = 12.5 events. 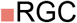, 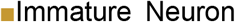, 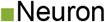, ■Zygote. (B) The same lineage in cladogram form with supporting nodal posterior probabilities ≥ 0.1 indicated. A “densiTree” representation of 1,500 sampled cladograms is shown as a mirror image. The complete densiTree representation is shown as an inset with the magnified area indicated by a black rectangle. ⬤ Mutations shared across members of a particular node; ▲ mutations that are shared in all but one member of a particular node. (C) Barcode of segregating alleles (*) from a monophyletic clade marked “C” in panel (B). Chromosomes are aligned with the centromere to the left. Blue and yellow regions indicate LOH events (B6 and CA, respectively). Green regions indicate heterozygosity. Node posterior probability is shown on the cladogram.

Posterior probability support for a specific topology is less robust. The consensus tree, depicted as a phylogram in **Fig. 3A** and a cladogram in **Fig. 3B**, is based on 4,502 sample trees, and the 99% credible set contains 4,457 trees. This is unsurprising, given the large polytomy at the base of the tree. A cladogram showing node support is shown in **Fig. 3B**, along with composite visualization of 1,500 overlaid trees from the credible tree set. For nodes with a posterior probability ≥ 0.1, the maximum level of support was 0.60 (median interquartile range). In general, resolved nodes correlate with the presence of at least one shared LOH event in daughter cells, and several clades share an allele in all but one cell (⬤ and ▲, respectively, along tree branches in **Fig. 3B**). For some clades, one allele appears sufficient to distinguish a clone, as shown by large dense wedges in **Fig. 3B**, but the topology of the monophyletic group cannot be resolved due to a lack of segregating alleles or confounding effects of coincidental identity by state (homoplasy). The informative chromosomal barcodes with segregating LOH alleles are shown for a representative clade in **Fig. 3C**. Taken together, these results indicate that there is a lineage signal driven by LOH events that is detectable using parsimony, but confounding variables such as homoplasy limit our ability to ascertain finer topological structure.

While we do not know the true relationship of the cells of any given mouse, pairwise analysis of any two mice can provide insight into the relationship between posterior probabilities and truly related cells, along with an estimate of homoplasy. Cells from a particular mouse should group together based on the principle of identity by descent while mixed groups should reflect homoplasy. An analysis of pooled cells from pairwise comparisons of mice, **Supplemental Fig. 6**, reveals that a posterior probability of 0.45 - 0.50 is likely sufficient for detecting clonal patches of cells.

### Stereotyped Expansion and Differentiation in Neocortical Histiogenesis

Histogenesis in the mouse neocortex consists of a progression of progenitor cells with decreasing proliferative potential (Taverna et al., 2014). Neuroepithelial stem cells divide symmetrically to produce the initial progenitor pool in the ventricular zone of the developing neocortex. This population transitions to an asymmetrical cell division stage consisting of RGCs to create unitary clusters of neurons either directly or through transient progenitors that expand the size of the cluster. A subset of RGC (about one-sixth) transitions to a gliogenesis state where they produce astrocytes and oligodendrocytes (Gao et al., 2014). Symmetrically dividing glial progenitors are produced as well (Ge et al., 2012). This pattern of symmetrical cell division followed by unitary asymmetrical cell division, expansion, and differentiation should produce multiple monophyletic clades of mixed cell types. We see such a pattern in the inferred phylogenies, **Fig. 4A**. In P0 mice, the diversity of cell types is limited, but monophyletic groups appear, consisting of neurons and RGCs. The P42 mice, which have increased cellular diversity, show the same topological pattern. Heterogeneous clades are predicted to be composed of neurons, immature and mature astrocytes, as well as oligodendrocytes and their intermediate progenitors, in agreement with a unitary expansion model of neurogenesis (Gao et al., 2014). Interestingly, the majority of glutamatergic neurons form their own clade in both P42 mice. In contrast, RGC and neurons in P0 mice are distributed throughout different clades. Correspondingly, we note that the large polytomies in each mouse consist of a similar number of monophyletic groups (15, 14, 14, and 11, for each of the four mice, respectively) though we cannot speak to whether or not this structure is meaningful, again due to low cellular sampling.

**Figure 4.**
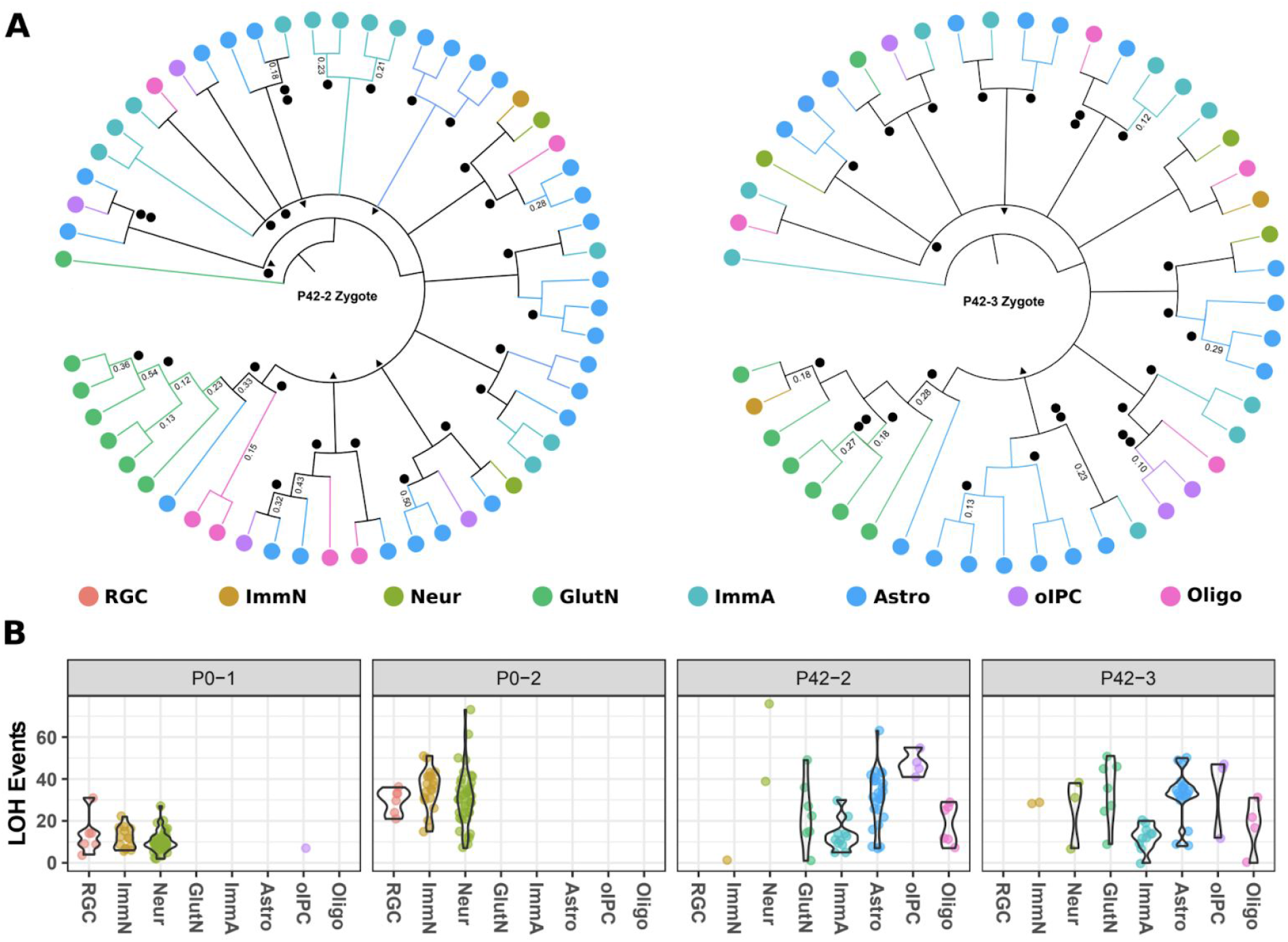
Stereotyped expansion in the mouse neocortex. (A) Consensus cladograms of two P42 mice. Nodes with posterior probability of >0.05 are resolved. Posterior probabilities of >0.10 are indicated. ⬤ LOH event shared among all daughter cells. ▲ LOH event shared in all but one daughter cell. (B) LOH events by mouse and cell type. Total autosomal LOH events are shown for each cell on the y-axis. Violin plots show the normalized density for each cell type in each mouse. RGC = radial glial cells, ImmN = immature neurons, Neur = neurons, GlutN = glutamatergic neurons, ImmA = immature astrocytes, Astro = astrocytes, oIPC = oligodendrocytic intermediate progenitor cells, Oligo = oligodendrocytes.

Generation of LOH events correlates with cell division (Mehta and Haber, 2014) and therefore likely reflects mitotic age. At P0, particularly for the P0-2 mouse, the distribution of the number of identified LOH events among RGC is tighter than seen in either group of neurons (**Fig. 4B**). The observation of fewer LOH events in more differentiated cells is consistent with RGC serving as a stem-like population, with neurons spun off earlier during embryogenesis. Conversely, the appearance of neurons exhibiting more LOH events is consistent with their production from older RGC populations along with subsequent additional cell divisions prior to terminal differentiation. At P42, the distribution of the number of LOH events in immature astrocytes is more tightly clustered than for mature astrocytes, and the median number of events is lower. Though we do not have contemporary RGC with which to compare, this pattern is consistent with immature astrocytes being produced from transitioning late-stage RGC, expanding, then differentiating into mature astrocytes. There appear to be at least two waves of oligodendrocyte production given that contemporary precursors are mitotically older than mature oligodendrocytes. Our findings are consistent with the concept of early vs. late populations of RGC producing neurons and glial cells, indicating that the differences we are observing in P42 mice are due to sampling from both populations (Kriegstein and Alvarez-Buylla, 2009).

## DISCUSSION

We have used scRNA-seq to simultaneously identify cell state based on gene expression profile and phylogenetically infer lineage by identifying LOH events corresponding to tracts of monoallelic expression.

The goal of our studies is to reconcile lineage with cell state trajectory. Corticogenesis in the mouse is complex and involves a linear transition of distinct progenitors that contribute to the production of neurons and macroglia at different stages of development (**Fig. 5A**). Due to the asymmetrical nature of RGC division and potential symmetrical expansion of some daughter cells, one RGC can contribute to multiple cell types and can tie cells produced at later stages of life to cells produced embryonically. Depending on the timing of LOH events during this process, several possible cell-type-specific phylogenetic topologies are possible. An event in neuroepithelial stem cells or early RGC would lead to mixed cell type clades while events occurring during the expansion of intermediate progenitor cells would mark more homogeneous clades. In P0 mice we observe mixed clades consisting of RGC, immature neurons, and neurons, a pattern that is highlighted for large clades when member cells are plotted in dimensionally reduced space based on transcription (**Fig. 5B**). These mixed clades are also seen in P42 mice (**Fig. 5C**), where cell types are more diverse given maturity. Combined, these results suggest that the large polytomies in all four mice might correspond to the initial expansion of neuroepithelial stem cells during embryogenesis and are consistent with stereotyped neuronal expansion. Interestingly, we do observe clades of predominately glutamatergic neurons in both P42 mice. This could be a reflection of neurons produced from earlier RGC or could be an artifact of relatively low cellular sample sizes compared to the total number of cells in the cortex. Increases in cellular sampling from the whole cortex or sampling from a much smaller region of cortex would allow greater resolution, and we would also expect to see increases in more cellularly uniform groups due to detection of events that occur during expansion of intermediate progenitor cells (Kriegstein and Alvarez-Buylla, 2009).

**Figure 5.**
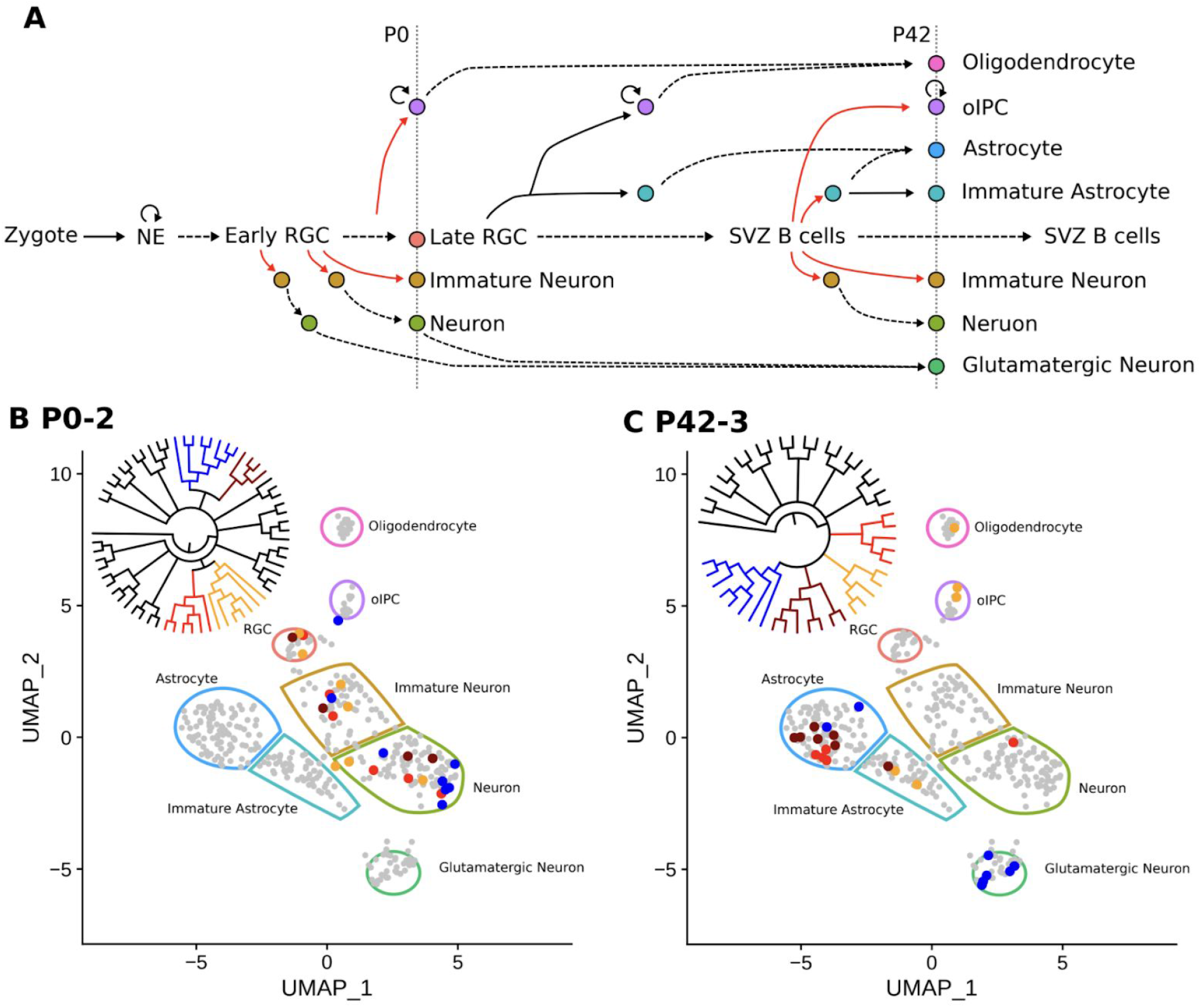
Lineage across cell types. (A) Expansion and differentiation of Emx1^+^ neural stem cells (NSC) into neurons and glia. Neuroendothelial (NE) cells expand (circular arrow) to form a pool of radial glial cells (RGC) that produce neurons and glia in the cortex via asymmetric cell division (red lines), expansion, and maturation (dashed lines). Time-dependent development proceeds horizontally. Observed cells in this study are indicated by vertical dotted lines along with their relative time points. oIPC = oligodendrocyte intermediate progenitor cell, SVZ = subventricular zone, as adapted (Kriegstein and Alvarez-Buylla, 2009). (B) Example of different clades encompassing multiple cell types for a P0 mouse. All 404 cells are shown in reduced dimensional (UMAP) space to illustrate all possible cell types, indicated by color-bound regions. Cells from four representative clades of the P0-2 mouse (inset cladogram) are indicated by their corresponding colors. (C) Same as in (B) for mouse P42-3.

As far as we are aware, our analysis provides the first indication that monoallelic expression observed in scRNA-seq datasets occurs in contiguous clusters unique to different cells, which we interpret as consistent with LOH.

Among inbred strains of mice, LOH ordinarily goes undetected in experiments employing scRNA-seq because the diploid genome is entirely homozygous. Here we examined F1 offspring of different mouse strains in order to generate frequent and predictable heterozygous SNVs distributed throughout the genome, focusing on transcribed loci, which can consequently be readily discriminated through scRNA-seq.

Different cell types express different genes, meaning that the same SNVs distinguishing monoallelic expression will not always be informative when comparing different cells. A challenge in defining contiguous stretches of monoallelic expression shared by different cell types therefore relates to identification of their boundaries when they occur within a gene expressed in a tissue specific manner. However, across a sufficiently large chromosomal region, at least several genes are likely to be expressed in any particular cell, allowing more precise restriction of boundaries and reducing the likelihood that homoplasy will confound phylogenetic reconstruction of lineage.

For any particular heterozygous SNV, there are several reasons why transcripts corresponding to only one parental allele may be detected, including limited sampling due to inadequate depth of coverage, capturing a moment in time when only one allele is actively undergoing transcription (i.e., “bursting” (Borel et al., 2015; Deng et al., 2014; Finn and Misteli, 2019; Reinius and Sandberg, 2015; Reinius et al., 2016)), and imprinting or other forms of allelic exclusion, such as X-inactivation (Galupa and Heard, 2018) or involving antigen receptors in the immune system or olfactory receptors (Khamlichi and Feil, 2018).

Regarding the latter, random mitotically stable monoallelic expression of some genes (Savova et al., 2016), such as *Pax5 (Nutt et al., 1999)* and *Il2 (Holländer et al., 1998)*, has been proposed as a source of diversification of somatic cells. However, these observations prove controversial. For example, monoallelic expression of *Nanog* has been posited to regulate pluripotency (Miyanari and Torres-Padilla, 2012), while other reports find that its expression is biallelic and no different from other genes playing similar developmental roles (Faddah et al., 2013; Filipczyk et al., 2013).

Reinius et al. (Reinius et al., 2016) addressed the frequency of monoallelic expression using scRNA-seq. In primary B6×CA mouse fibroblasts, they found that 13% of autosomal genes exhibited monoallelic expression. To determine if monoallelic expression corresponded to a clonal distribution, scRNA-seq data were pooled from fibroblasts clonally expanded *in vitro*, at which point monoallelic expression was observed in no more than 0.5% of expressed genes and was therefore interpreted as dynamic and consistent with bursting. It is worth noting, however, that they excluded genes from regions with cell- or clone-specific chromosomal aberrations, which might underlie the clonal abnormalities we infer to represent LOH events.

In further elegant studies, Reinius et al. isolated antigen-specific T cells following yellow fever virus vaccination in a human subject, under the premise that “rearrangements of the two T cell receptor (TCR) chains result in immense sequence variability [such that] cells with identical rearrangements can be identified as clones”. Counterintuitively, however, multiple individuals frequently share an identical, “public” TCR responding to the same antigen, due to biases and convergence in TCR rearrangements (Elhanati et al., 2018; Li et al., 2012; Madi et al., 2014; Pogorelyy et al., 2018a). In fact, in identical twins, about one-third of T cells responding to Yellow Fever vaccination share TCR sequences, which is concluded to arise from multiple independent clones undergoing convergent recombination and selection (Pogorelyy et al., 2018b). Therefore, T cells with identical TCR rearrangement may frequently have polyclonal origins. An additional limitation is that Reinius et al. restricted their scRNA-seq analysis to just ~800 autosomal genes. Finally, it is worth emphasizing that they nevertheless detected clonal monoallelic expression at about 1% of loci, with a range reaching much larger values among some T cells.

As an alternative to evaluating allele-specific polymorphisms, Nag et al. (Nag et al., 2013, 2015), sought a characteristic chromatin signature simultaneously exhibiting marks associated with active transcription (H3K36me3) and silencing (H3K27me3) and predicted that monoallelic expression occurs for ∼20% of ubiquitously expressed genes and over 30% of tissue-specific genes across cell types.

In contrast, for all cells of all four mice analyzed for LOH, we observed a mean of 5.1% (range 0 - 19.1%) of the autosomal genome exhibiting monoallelic expression, a result much closer to Reinius et al. than to Nag et al. One reason the value we determined may exceed Reinius et al. is that our analysis of heterogeneous cell types required binning of the boundaries of regions of monoallelic expression to account for differences in gene expression between individual cells, which necessarily leads to overestimation.

We emphasize that for any given locus exhibiting monoallelic expression, we infer that the vast majority of cells, including from the same cell type, exhibit a heterozygous genotype, which would not be true if there were pervasive random clonal monoallelic expression. We agree with the conclusion of Reinius et al. that random clonal monoallelic expression is an infrequent gene regulatory mechanism and is therefore unlikely to confound interpretation of LOH events. To further control for this issue, we surveyed for large, contiguous tracts of monoallelic expression, spanning multiple genes, beyond distances generally correlated with transcriptional bursting or topologically associated domains, thereby attempting to minimize artifactual detection of monoallelic expression. To assure that segmental clustering of monoallelic expression was similarly not a random phenomenon or an artifact of the algorithm employed for its detection, we found that the tracts largely disappeared when SNV genotypes were permuted and that shared LOH events were enriched in cells from the same, as opposed to different, mice.

Several recombinational pathways can lead to LOH, whether it be copy neutral or result in loss of chromosomal material (Mehta and Haber, 2014). Analysis of these pathways at a single cell level may be helpful in defining LOH events in cancer and elucidating how some heritable disorders undergo somatic reversion (Revy et al., 2019), in addition to its application in developmental biology.

A mechanism of LOH consistent with our observations is gene conversion, resulting in copy neutral LOH. The size and frequency of LOH we detect is comparable to prior observations. Nevertheless, we cannot exclude LOH arising due to deletion of the undetected allele. While the density of informative SNVs in our experiments is comparable to SNV microarrays clinically employed for detection of copy number variants, cell-to-cell differences in gene expression level would prove challenging for purposes of copy number analysis.

We cannot entirely exclude epigenetic silencing of one allele, as proposed as an explanation for monoallelic expression (Andergassen et al., 2017; Nag et al., 2013, 2015). However, for that to occur would somewhat implausibly require coordinated epigenetic regulation of multiple adjacent genes, including across different cell types. Moreover, the occasional observation of single cell biallelic expression at some loci on the X chromosome or the PWS/AS region engender confidence in our calls of LOH, as these “escapes,” or otherwise apparently leaky expression, indicate that an underlying heterozygous genotype remains determinable when large regions are epigenetically silenced. Even if epigenetic phenomena were contributing to our calls of LOH, the information would likely still prove useful for inferring lineage. In fact, an early approach for retrospectively inferring lineage from sequencing data employed detection of cytosine methylation, involved in developmentally regulated gene silencing (Shibata and Tavaré, 2007).

The number of possible lineage histories grows explosively as a function of the number of cells (Salipante and Horwitz, 2007), complicating phylogenetic approaches for inferring lineage. A workaround to this issue could come in the form of a deterministic barcode, similar to a “blockchain” (Ozercan et al., 2018), in which the sequence of cell divisions and daughter cell relationships to one another were explicitly recorded. In such a situation, the number of barcodes required to unambiguously reconstruct the cell lineage would simply equal the number of cells. Intriguingly, mitotic recombination is typically reciprocal, such that heterozygosity at a locus undergoing recombination is expected to be inherited by one daughter cell as homozygosity for one parental allele and, for the other daughter cell, as homozygosity for the opposite parental allele. It is possible that by performing scRNA-seq on a larger number of cells, reciprocal copy neutral LOH outcomes of a single recombination event could be identified and lend greater certainty to lineage reconstructions.

A distinct advantage to the approach described here, compared to recently introduced technologies for dynamically barcoding cells through genome editing, is that it can be performed retrospectively, permitting its use for studying development in humans and other organisms where genetic engineering or other forms of embryonic manipulation are infeasible or where lifetimes are long. Unlike laboratory mouse strains, humans and individuals from other species with large populations are outbred, and heterozygous SNVs are abundant, making the method applicable without any special breeding strategy, provided that the underlying phased genome sequence is determinable using experimental and/or computational approaches (Choi et al., 2018). For example, our approach may prove particularly informative in evaluating the lineage of the earliest events in cancer, where there may be fewer mutations to otherwise infer clonality, or in studying aging in long-lived organisms, where it is most convenient to sample endpoints late in life.

Because LOH events are likely to be cumulative throughout the lifespan of an organism, an additional source of information for inferring lineage relates to the total number of LOH events. In contrast, some of the genetically engineered barcoding strategies lead to a concentration of mutations at early stages of embryogenesis. Combining both approaches could prove complementary and help improve the reliability of sequence-based mass scale lineage reconstructions.

A limitation of our approach is that it requires both a breadth and depth of scRNA-seq sufficient to infer the allelic origin of SNVs contained within a transcript, compared to more limited sampling necessary to simply identify a transcript’s cognate gene. As throughput increases and nucleic acid sequencing costs continue to decline, this may become less of an issue.

## METHODS

### Mice

The dataset we used for our analysis was previously published; mouse strains, breeding strategy, cell isolation, and scRNA-sequencing approaches are as described (Laukoter et al., 2020). Briefly, B6 (*Emx1*-Cre;*Z/EG*) and CA mouse strains obtained from The Jackson Laboratory were bred and analyzed in accordance with protocols approved by Institutional Animal Care and Use Committees at IST Austria. Cells were dissociated from cerebral cortex, and *Emx1*^+^ single cells were sorted by FACS for library preparation. RNA sequencing data can be accessed using the GEO Series accession number GSE152716 (http://www.ncbi.nlm.nih.gov/geo/query/acc.cgi?acc=GSE152716).

### scRNA-seq

RNAseq cDNA libraries were prepared from single cells using Smart-seq2 (Picelli et al., 2013). Single-end 50 base pair (bp) reads were mapped to GRCm38.p5 (mm10) and expression determined using Ensmbl [91, Dec 2017] with STAR 2.5.0c (Dobin et al., 2013), as previously described (Laukoter et al., 2020). A million or greater total reads and a range of 10,000-30,000 total mRNAs were used to filter high quality single cell samples. 1,735,775 unique reads were mapped to the median cell (Lower Quantile (LQ) 990,582; Upper Quantile (UQ) 2,858,522).

### Cell Identification

Cell identity was determined through the Seurat R package (Butler et al., 2018; Stuart et al., 2019). Gene level expression data from all 404 cells from 8 mice were combined and filtered. Genes expressed in a minimum of three cells were analyzed. Cells with a minimum of 200 genes detected were kept for further inspection. All 404 cells passed the criteria above resulting in a 404 × 23,373 feature count matrix (**Supplemental Data**). The count matrix was normalized with the NormalizeData function, setting normalization.method to “LogNormalize” and scale.factor to 10,000. Each cell was assigned a cell cycle state using the CellCycleScoring function. The list of S and G2/M genes is provided in **Supplemental Table 2**. The expression matrix was scaled (mean expression = 0, variance = 1) so that highly-expressed genes do not dominate downstream analysis. We also removed variation due to mitochondrial contamination, library preparation (individual mouse), mouse age, and cell cycle stage (S or G2/M) using the vars.to.regress option in the ScaleData function. We reduced the dimensionality of the expression data via principal component analysis (PCA) of the top 2,000 variably expressed genes (selection.method = “vst”) and UMAP, using the first 15 components. Cells were clustered using Seurat’s graph-based clustering tools. A K-nearest neighbor graph, based on Euclidean distance in PCA space, was constructed for all cells using the FindNeighbors function. The Louvain algorithm (FindClusters(resolution = 0.8)) was then used to iteratively group cells together. Clustered cells were then annotated based on marker gene expression patterns (**Supplemental Table 1**).

### scRNA-seq Variant Calling

Further studies focused on a subset of four mice (P0-1, 56 cells; P0-2, 64 cells; P42-2, 56 cells; and P42-3, 47 cells). Heterozygous sites were identified using the CA Mouse Genome Project sequencing data (CAST_EiJ.mgp.v5.dbSNP142.vcf) (Keane et al., 2011). Briefly, homozygous SNV loci for CA, when compared to mm10 (B6), passing GATK hard filtering metrics for SNV detection based on DNA sequencing were identified and used as a guide (ROD file) for loci to be analyzed in GATK. 20,667,142 SNV loci were identified. On average, this provides 8,392 loci per Mb coverage of autosomes and X chromosomes. RNA-based SNV calls were made at predicted sites of heterozygosity in B6:CA F1 mice using the GATK (Auwera et al., 2013) variant calling tool (-T GenotypeGVCFs -stand_call_conf 20 -stand_emit_conf 10) and hard filtered using recommended settings for scRNA-seq variants (-T VariantFiltration -window 35 -cluster 3 -filterName FS -filter “FS > 30.0” -filterName QD -filter “QD < 2.0”).

### Genotype Determination

A cell’s genotype at a particular locus was inferred by modeling the scRNA-seq variant data as a hidden Markov process with the underlying genotype at a particular locus corresponding to the hidden state and the scRNA-seq-based variant status as the emission or observation state.

The hidden Markov model (HMM) contains three hidden states and three emission states, described below.

The hidden genotype has three possible states: homozygous B6, heterozygous, or homozygous CA. The transition matrix between states is defined as:

|  | Homozygous B6 | Heterozygous | Homozygous CA |
| --- | --- | --- | --- |
| Homozygous B6 | 0.99999 | 0.00001 | 0 |
| Heterozygous | 0.00010 | 0.99980 | 0.00010 |
| Homozygous CA | 0 | 0.00001 | 0.99999 |

The observed scRNA-seq-based genotype (monoallelic B6, biallelic B6:CA, or monoallelic CA) is based on the emission state probability of the underlying hidden state given here:

| Hidden\Emission state | Monoallelic B6 | Biallelic | Monoallelic CA |
| --- | --- | --- | --- |
| Homozygous B6 | 1 | 0 | 0 |
| Heterozygous | 0.45 | 0.10 | 0.45 |
| Homozygous CA | 0 | 0 | 1 |

The Markov-chain’s starting state probability is given as 0.05 for each of the homozygous states and as 0.90 for the heterozygous state.

For any given chromosome of any given cell the underlying genotypes of scRNA-seq detected loci were determined by the most probable order of hidden states (Viterbi path) based on the scRNA-seq variant data using the Viterbi algorithm contained in the HMM R package (https://cran.r-project.org/web/packages/HMM/index.html). The Viterbi path describes the amalgamation of both homologous chromosomes and is assumed to have a copy number of two. Hemizygosity will appear as LOH, and duplications will appear as heterozygous (**Supplemental Data**).

### Identification of LOH Events

A single transition from the hidden heterozygous state to either homozygous state could indicate recombination between two homologous chromosomes that extends to the telomere. A subsequent transition from the homozygous state back to the heterozygous state would describe an interstitial event such as a double crossover or non-crossover strand invasion. Partial or micro chromosomal deletions would also trigger these transitions. Transcriptional bursting kinetics or epigenetic silencing on one allele could account for short runs of either B6 or CA alleles, as well, confounding our results. To control for such events we removed homozygous runs that spanned a region ≤1 Mb.

Due to non-uniform scRNA-seq coverage across cells we grouped LOH events in 2 Mb bins, starting at the beginning of each chromosome. Events were characterized by their beginning and ending bins along with their parental allele identity.

### Permutation Analysis

For each mouse, a subset of randomly sampled “cells” was created *in silico* based on the observed scRNA-seq variants. For each locus, an scRNA-seq allele state (no coverage, biallelic, monoallelic B6, or monoallelic CA) was randomly sampled with replacement from the pool of allele states at that locus across all cells from the mouse in question. This process was repeated for all analyzed loci along the length of each autosome. These virtual cells were then genotyped as described above, and LOH events were identified for each generated cell (**Supplemental Data**).

### Encoding LOH Events for Cladogram Construction

Tracts of LOH >1 Mb are described by four elements: their chromosome location, the 2 Mb bin in which they start, the strain identity of the tract (B6 or CA), and their ending 2 Mb bin. For example, a homozygous run on chromosome 19, starting at position 5,000,000 and ending at position 9,000,000, and consisting exclusively of CA variants would be described as chr19_3CA5. A chromosome with no LOH event is coded as HT, designating that the chromosome as apparently heterozygous (e.g., for chromosome 19 the designation would be chr19_HT). A chromosome for any one cell can contain multiple LOH events.

Each chromosome is considered its own independently evolving region within a mouse, with the cell specific combination of LOH events defining its state. An LOH event occurring over a specific region of the chromosome does not preclude another event happening nor does it affect the probability of that event happening. Once a heterozygous stretch of chromosome is converted to a homozygous run it cannot revert back to the homozygous state. For any chromosome, each LOH event is treated as independent and irreversible. The multiple character states for any chromosome in any mouse can be factored into binary characters representing the presence (1) or absence (0) of any chromosome specific LOH event. For example, consider the state of chromosome 19 in three fictitious cells:

Cell01: chr19_HT

Cell02: chr19_2B65, chr19_20CA30

Cell03: chr19_2B65, chr19_15B626

There are three unique LOH events with one cell lacking any event. These cells would be factored and encoded as:

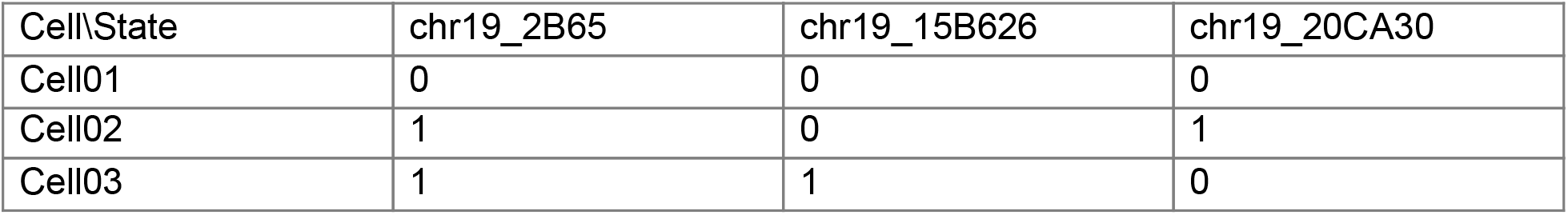

| Cell\State | chr19_2B65 | chr19_15B626 | chr19_20CA30 |
| --- | --- | --- | --- |
| Cell01 | 0 | 0 | 0 |
| Cell02 | 1 | 0 | 1 |
| Cell03 | 1 | 1 | 0 |

### LOH Evolution Model

We implement a Camin-Sokal (Camin and Sokal, 1965) parsimony-inspired Bayesian model of LOH evolution to infer lineage relationships, using the restriction site model in the MrBayes software package (Ronquist et al., 2012). It is assumed that the zygote for any F1 mouse is biallelic across the assayed SNVs for any particular autosome (a state of 0 in the coding scheme described above) and that this is the ancestral condition. An LOH event creates a new character state (1), and the transition from 0 to 1 is 100-fold more likely than the reverse. Though LOH events can occur between two X-chromosomes, sex chromosomes are not considered here because two mice are male and X-inactivation in female mice changes assumptions of the HMM used to infer genotypes based on RNA. Visualization of the Bayesian posterior distribution of cladograms employs the DensiTree algorithm (Bouckaert, 2010).

## ACKNOWLEDGEMENTS

We thank Bill Bolosky, Microsoft Research, for earlier work showing proof of concept in TCGA bulk RNA-seq data. Supported by the Paul G. Allen Frontiers Group (University of Washington); NIH R00HG010152 (Dartmouth); and NÖ Forschung und Bildung n[f+b] life science call grant (C13-002) to SH, and the European Research Council (ERC) under the European Union’s Horizon 2020 research and innovation program 725780 LinPro to SH.

## AUTHOR CONTRIBUTIONS

All authors conceived the experiments. FMP and SH performed mouse experiments and scRNA-seq analysis. DJA and MSH perform LOH and phylogenetic analysis. DJA and MSH drafted the manuscript with contributions from all other authors.

## SUPPLEMENTAL MATERIALS

**Supplemental Figures (appended below)**

**Supplemental Table 1. Genes used to identify cell types**, xlsx file.

**Supplemental Table 2. Cell cycle related genes**, xlsx file.

**Supplemental Table 3. Genes escaping X-inactivation**, xlsx file.

**Supplemental Table 4. LOH allele enrichment**, xlsx file.

**Supplemental Data (ZIP archive):** https://drive.google.com/file/d/1gupKbsqf5mLj_b3OeIqlrihFxP2_eurh/view?usp=sharing

**Seurat_CountMatrix.txt**

The count matrix of gene level transcription from 404 cells used in combination with Seurat. 23,373 features.

**Folder - Consensus_Trees**

“Majority rule” consensus trees for each mouse and mouse comparisons in NEXUS format.

**Folder - HMM_Genotype_Tables**

scRNA-seq based HMM genotype calls for all 19 autosomes. This folder contains two sub-folders. HMM_Genotype_Tables_Mouse contains the TSV files for P0-1, P0-2, P42-2, and P42-3. HMM_Genotype_Tables_Sampled_10K contains the tar.gz compressed TSV files based on the in silico sampled 10,000 “cells” for P0-1, P0-2, P42-2, and P42-3.

---

**Supplemental Figure 1.**
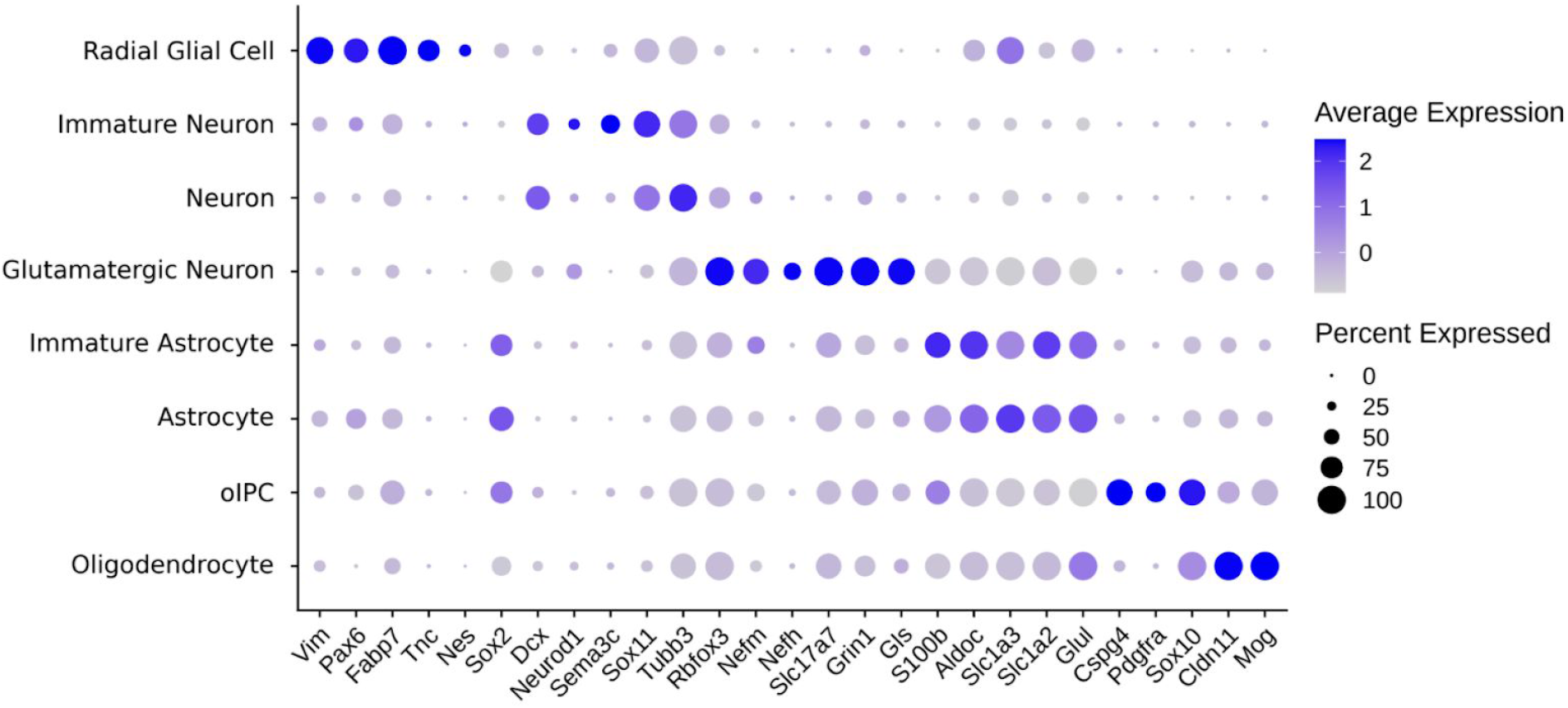
Mean cluster expression of cortical cell marker genes. Scaled average cluster expression is shown by color for each gene. Percent of cells in each cluster expressing the gene is indicated by dot size. Radial glial markers: *Vim*, *Pax6*, *Fabp7*, *Tnc*, *Nes*, *Sox2*, *Slc1a3*. Neuronal markers: *Dcx*, *Neurod1*, *Tubb3*, *Rbfox3*, *Nefm*, *Nefh*, *Slc17a7*, *Grin1*,*Gls*. Migration markers: *Sema3c* and *Sox11*. Astrocyte markers: *S100b*, *Aldoc*, *Slc1a3*, *Slc1a2*, *Glul*. Oligodendrocyte markers: *Cspg4*, *Pdgfra*, *Sox10*, *Cldn11*, *Mog*. oIPC, oligodendrocyte intermediate progenitor cell.

**Supplemental Figure 2.**
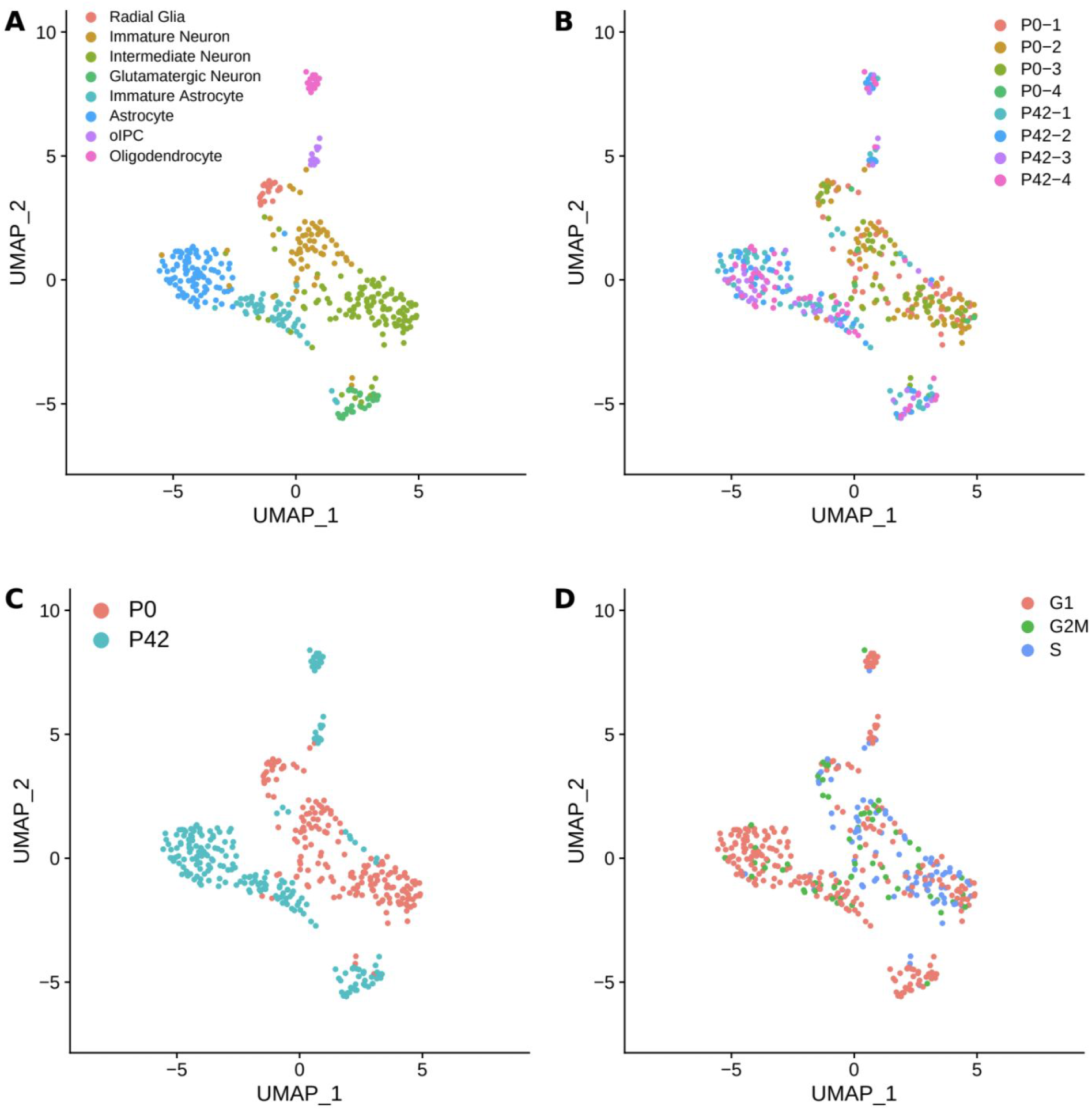
Distribution of mouse identity, age, and cell cycle state across individual cells. 404 neocortical cells plotted in dimensionally reduced (UMAP) space based on gene expression. Individual cells (dots) are colored based on (A) cell type (repeated here for convenience from Fig. 1B), (B) mouse identity, (C) age, and (D) cell cycle phase determined by expression of cell cycle related genes.

**Supplemental Figure 3.**
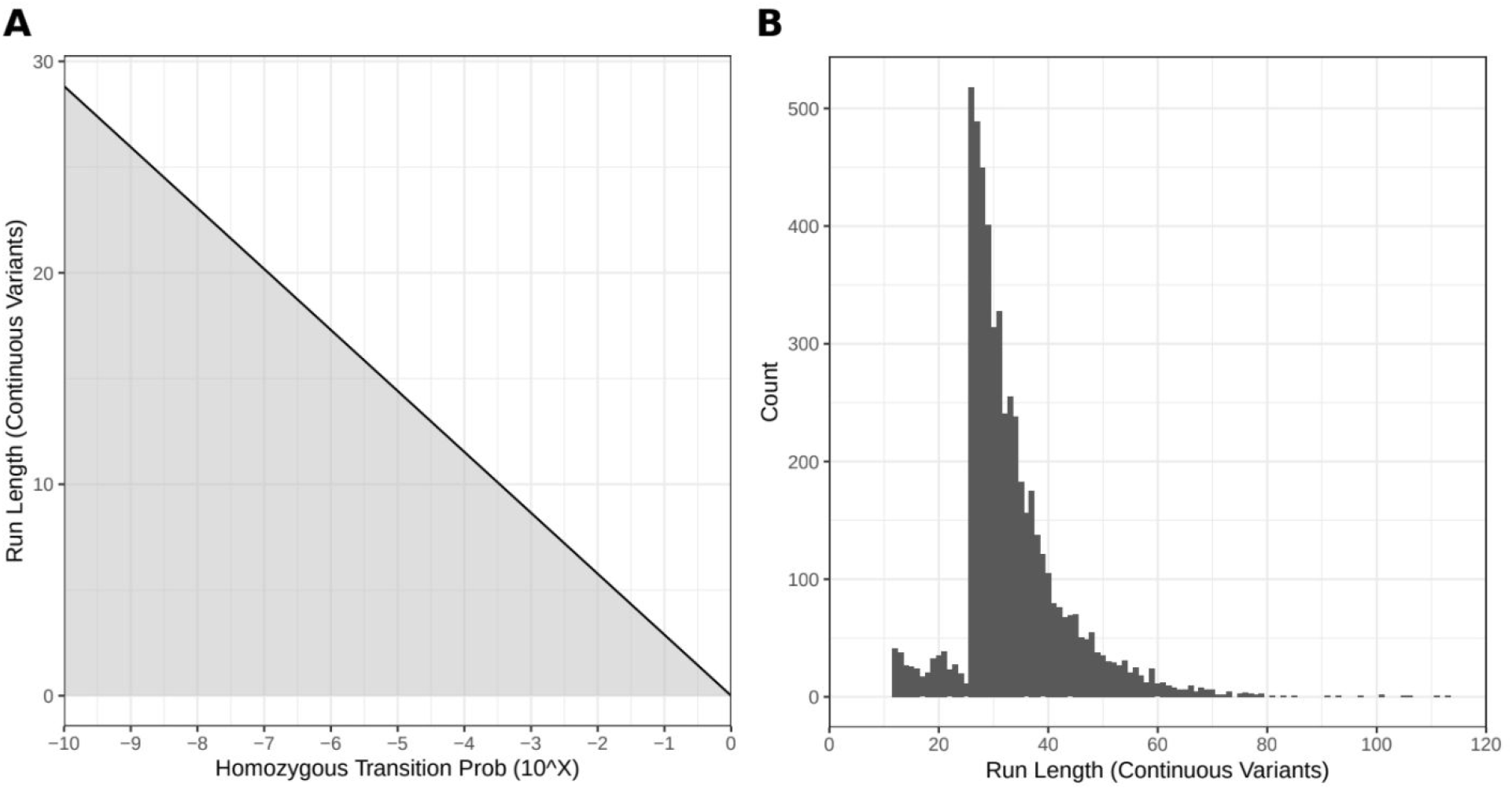
The required number of continuous monoallelically expressed variants for a predicted LOH event, as shaped by the homozygous state transition probability and the emission matrix. (A) The number of continuous SNV loci observed as monoallelic needed to trigger a homozygous hidden state change is shown as a function (black line) of the homozygous transition state probability (given the emission matrix described in the methods section). The gray region indicates the area where the hidden state remains heterozygous while the white region describes the conditions needed to trigger a hidden state change to homozygous for either allele, thus indicating an LOH event. (B) A histogram showing consecutive variant loci with length of LOH events ≥1 Mb for all four mice analyzed. A predominant number of LOH events contain at least 26 consecutive variants. These events would still be called even if the homozygous transition probability were 10^−8^.

**Supplemental Figure 4.**
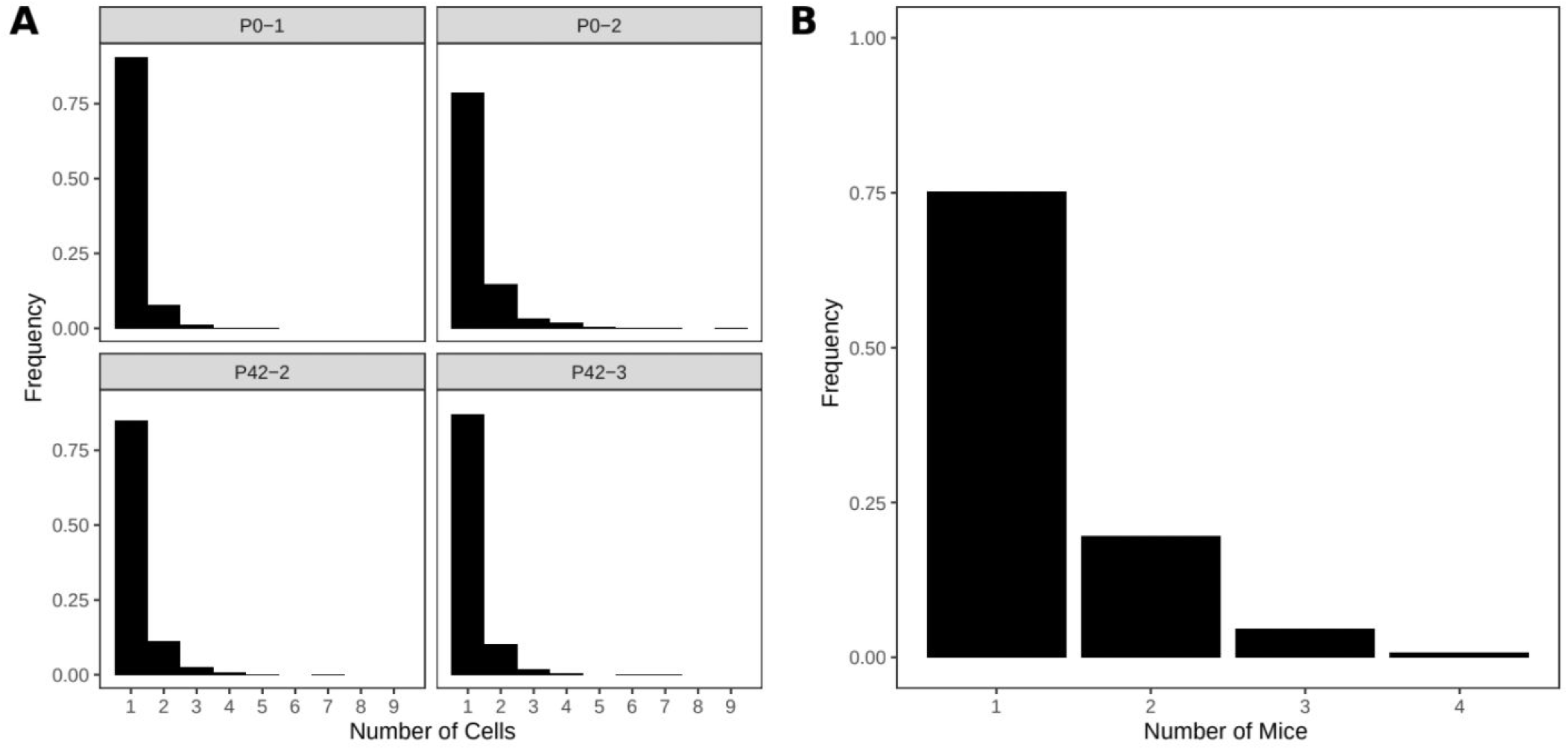
Distribution of shared alleles within and across mice. (A) Frequency of shared LOH events (alleles) across cells within an individual mouse. Total number of alleles for each mouse; P0-1 = 557, P0-2 = 1544, P42-2 = 1243, P42-3 = 1080. (B) Frequency of shared LOH events (alleles) across four mice. Total number of alleles = 3380.

**Supplemental Figure 5.**
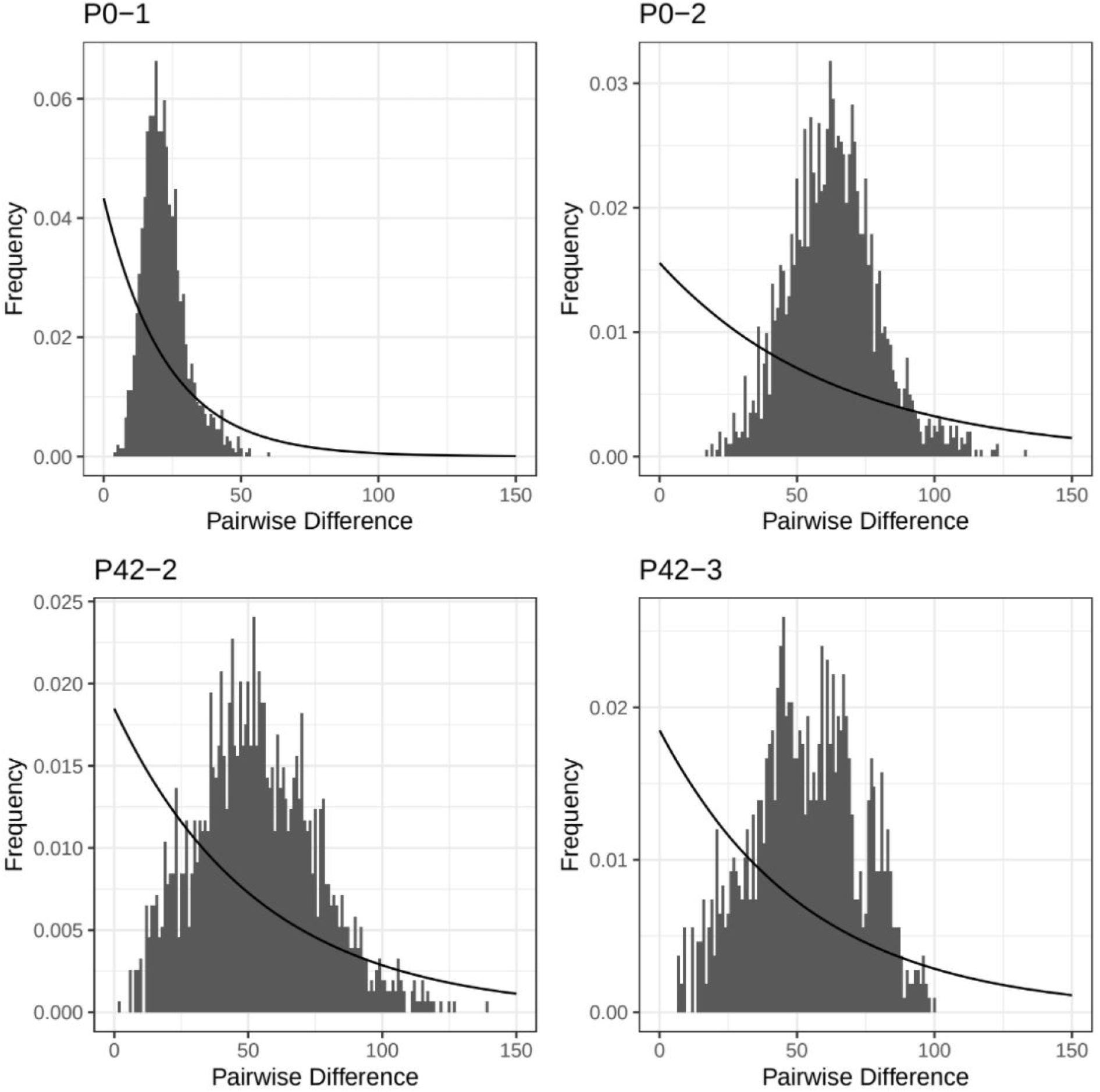
Distribution of pairwise differences between individual cells. For each mouse the distribution of pairwise differences between individual cells is shown. The solid line represents the expected distribution of differences in a population of constant size given the average pairwise distance for the sample. Populations undergoing exponential growth are predicted to have unimodal distributions of pairwise differences.

**Supplemental Figure 6.**
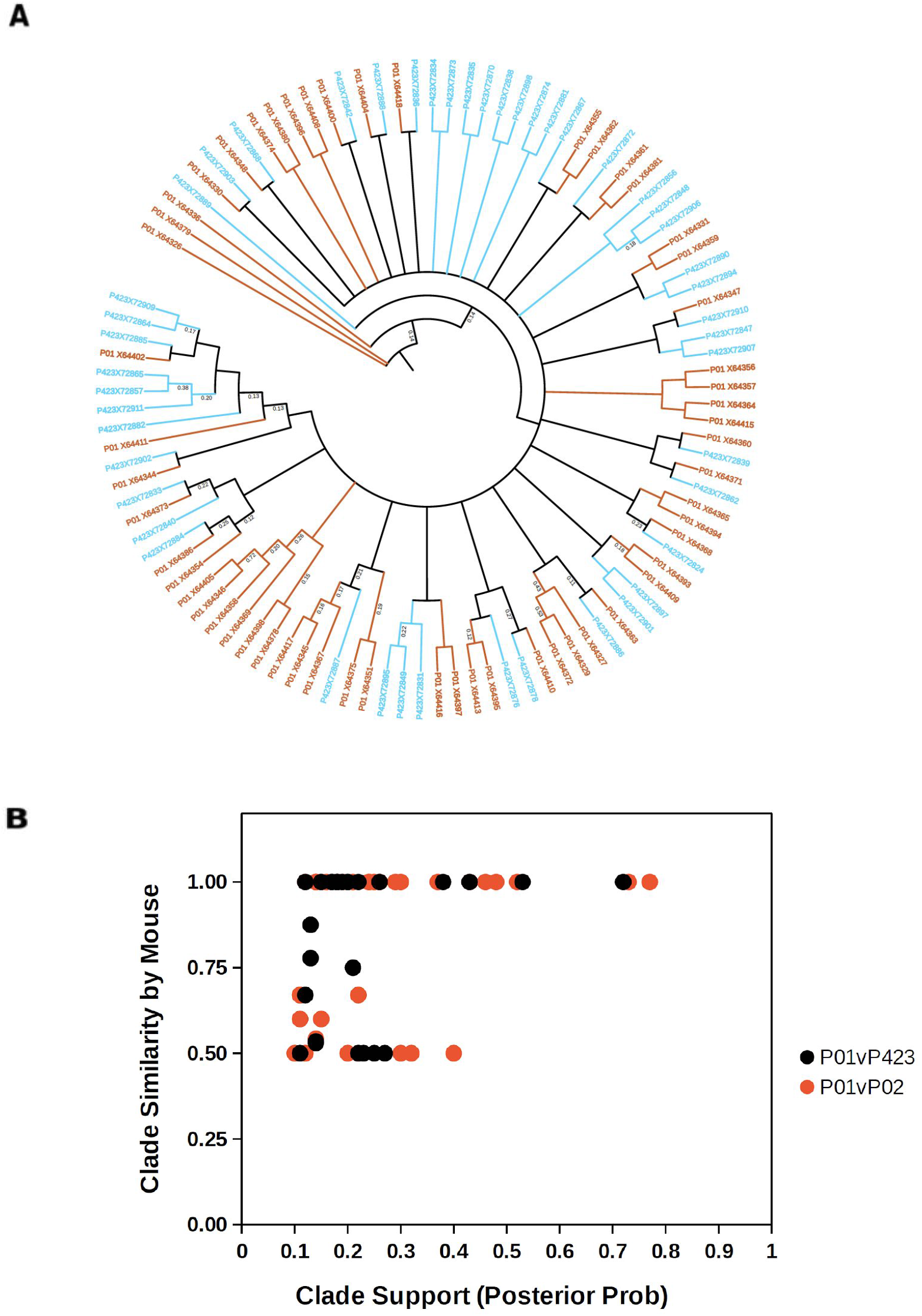
Clade support derivation from pairwise mouse mixing. (A) Phylogenetic analysis of two mice used to estimate meaningful posterior probability values, orange = mouse P0-1, blue = mouse P42-3. (B) Scatter plot of clade similarity (mouse identity) and posterior probability of clade suggesting a value of 0.45 - 0.50 as meaningful clonal designation, even if topology is not fully resolved.

## Notes

https://drive.google.com/file/d/1gupKbsqf5mLj_b3OeIqlrihFxP2_eurh/view?usp=sharing

### Competing Interest Statement

The authors have declared no competing interest.

